# Is it possible to keep the exoskeleton of the crab *Callinectes ornatus* soft for several days?

**DOI:** 10.1101/156638

**Authors:** Diogo Barbalho Hungria, Ubiratã de Assis Teixeira da Silva, Leandro Ângelo Pereira, Ariana Cella-Ribeiro, Antonio Ostrensky

## Abstract

Soft-shell crab is considered a gastronomic delicacy, reaching high values in the international market. The process of hardening of the crab’s exoskeleton after moulting takes approximately two days to complete; however, the duration for which the shell remains at the consistency of high commercial value is only 3 hours on average. After this period, the shell assumes a consistency classified as “paper”, later becoming “hard” again. The goal of this work was to evaluate the use of the crabs themselves to alter the chemical characteristics of the water and thereby increase the amount of time during which they can be marketed as “soft-shell crab”. In this work, 241 individuals of *Callinectes ornatus* were used in two experiments. In the first experiment, the animals were maintained in a collective system with filtration and partial daily water renewal. In the second experiment, the crabs were maintained in a collective system with filtration but no water renewal. In Experiment 1, the chemical characteristics of the water remained unchanged over time (p > 0.05), and the median time to hardening of the exoskeleton to the paper consistency after moulting was 3 hours. Over the course of Experiment 2, there was a significant reduction (p < 0.05) in pH and significant increases in the ammonia and nitrite concentrations. When moulting occurred in water with a pH below 7.3 and total ammonia concentrations above 6.0 mg/L, the crabs’ shells did not harden, and it was possible to keep them soft for up to 5 days.

## Introduction

*Callinectes ornatus* Ordway, 1863 (Crustacea, Decapoda, Portunidae) is a swimmer crab found from North Carolina (USA) to the Rio Grande do Sul (Brazil). It occurs in areas with sand, mud or shell bottoms and inhabits estuarine to marine areas at a depth of approximately 75 m (Carvalho and Couto, 2011; Melo-Filho, 1996).

Similar to other arthropods, *C. ornatus* grows through a process of periodic exoskeleton changes; each shedding of the exoskeleton is known as ecdysis or moult (Drach, 1939; Freeman and Perry, 1985; Newcombe et al., 1949). Immediately after shedding its exoskeleton, the crab presents a soft and flexible integument that has a low level of calcification. In this phase, the animals can be commercialised and consumed whole as soft-shell crab, a delicacy that is appreciated worldwide and that reaches high market values (Gaudé and Anderson, 2011; Oesterling, 1995; Perry et al., 2010). According to FAO (2013), the annual revenue generated from the production and marketing of soft-shell crab in 2012 was more than US$ 940 million.

Immediately after moult, CaCO3 deposition begins on the protein matrix of the new exoskeleton. This process involves a complex system of absorption of Ca^2+^, CO2, and HCO3^-^ and the synthesis of CaCO3 and other elements (Greenaway, 1985; Perry et al., 2001; Wheatly, 1999; Zanotto and Wheatly, 2002). The initially fragile exoskeleton undergoes rapid hardening, providing rigidity and mechanical protection for the animal. Under natural conditions, the hardening of the exoskeleton takes about two days to complete (Cameron and Wood, 1985). During the hardening process, the exoskeleton can be classified into four sequential levels of consistency: soft, leather, paper and hard (Freeman et al., 1987). Only the first two are valued in the international market of soft-shell crabs (Gaudé and Anderson, 2011; Oesterling, 1995; Perry et al., 2010). However, the combined duration of the soft and leathery stages is very short in nature, rarely lasting more than 3 h (Cameron and Wood, 1985), which obliges commercial producers to inspect all of the animals stocked in the pre-moulting phase every 4 h on average (Oesterling, 1995). Extending the duration in which the crab shells remain at the consistencies of high market value would significantly reduce production costs (Perry et al., 2001). Furthermore, it would minimise the damage caused by rapid exoskeleton hardening, providing better quality and uniformity regarding the softness of the product.

The goal of this work was to test the viability of using the crab C. ornatus to alter the chemical characteristics of the water to extend the time during which the animals could be marketed as soft-shell crab.

## Material and Methods

### Crab collection and maintenance

Specimens of *C. ornatus* were obtained via trawling by professional fishers at the balneary of Shangri-la, municipality of Pontal do Paraná (25°37'S/48°25'O), Paraná, Brazil. Shrimp trawls 12 m in length and with 20 mm mesh were used. In each sampling campaign, on average, three trawls of approximately 50 min each were made. Immediately after crab collection from the net, the crabs were separated and transferred to two 70 L polyethylene tanks with lids, each containing 20 L of seawater. The tanks received continuous aeration supplied via an 18 W air compressor. Inside each tank were plastic screens with 2 mm mesh positioned to reduce contact and prevent fights between the animals and minimise injuries and deaths. Thereafter, 100% of the water was renewed every half hour during the campaign.

Immediately after capture, the animals were transported to the Marine Aquaculture and Restocking Center (CAMAR) of the Integrated Group of Aquaculture and Environmental Studies (GIA), Federal University of Paraná (UFPR), at Pontal do Paraná (25°41'29.94"S, 48°27'57.09"W). The time elapsed between animal capture and arrival at the laboratory was consistently less than 4 h. Animals that were not used were returned to the sea.

In the laboratory, the crabs were maintained in 1,000 L tanks containing 100 L of seawater (30 psu) supplied with constant aeration for approximately 6 h. This period was purposely short since a large proportion of the captured individuals were very close to moult. Dead animals were discarded, and the live animals were classified by sex. Then, the crabs were inspected to determine the phase of the moulting cycle. Those individuals at the pre-ecdysis phase were selected for the experiments based on macroscopic indicators (visualisation of an inner line along the edges of the fifth pair of pleopods) (Drach, 1939; Drach and Tchernigovtzeff, 1967; Wehrtmann and Mena-Castañeda, 2003). The selected individuals were weighed on an analytical balance (Marte AL 500c, Brazil; accuracy of 0.01 g) and measured (width of the carapace, measured as the distance between the base of the largest lateral spines) with a pachymeter.

### Pilot experiments

Two pilot experiments were carried out. The first experiment tested the influence of fasting on animal survival under laboratory conditions. The animals only began to die after 50 days without access to food. Based on this result and to potential feeding effects on water quality or the process of moult and hardening, the crabs were not fed during the 12 days of each of the main experiments.

The second experiment tested the influence of the non-renewal of water on hardening time. The time elapsed between moult and shell hardening was significantly higher under water non-renewal than under the periodic renewal of water. In addition, a higher frequency of moult was observed at night (between 18:00 and 06:00); this information informed the design of the experimental methodology described below.

### Experimental Design

In both experiments, the saltwater had been previously chlorinated and maintained under constant aeration for 24 h. After this period, residual chlorine was neutralised (with 50% sodium thiosulfate), and the water was stored in the dark in 25,000 L tanks. Before use in the experiments, the water was passed through mechanical filters of 5 and 25 μm mesh and a UV filter for disinfection. Two experiments were performed and are represented schematically in Figure 1.

**Figure 1.**
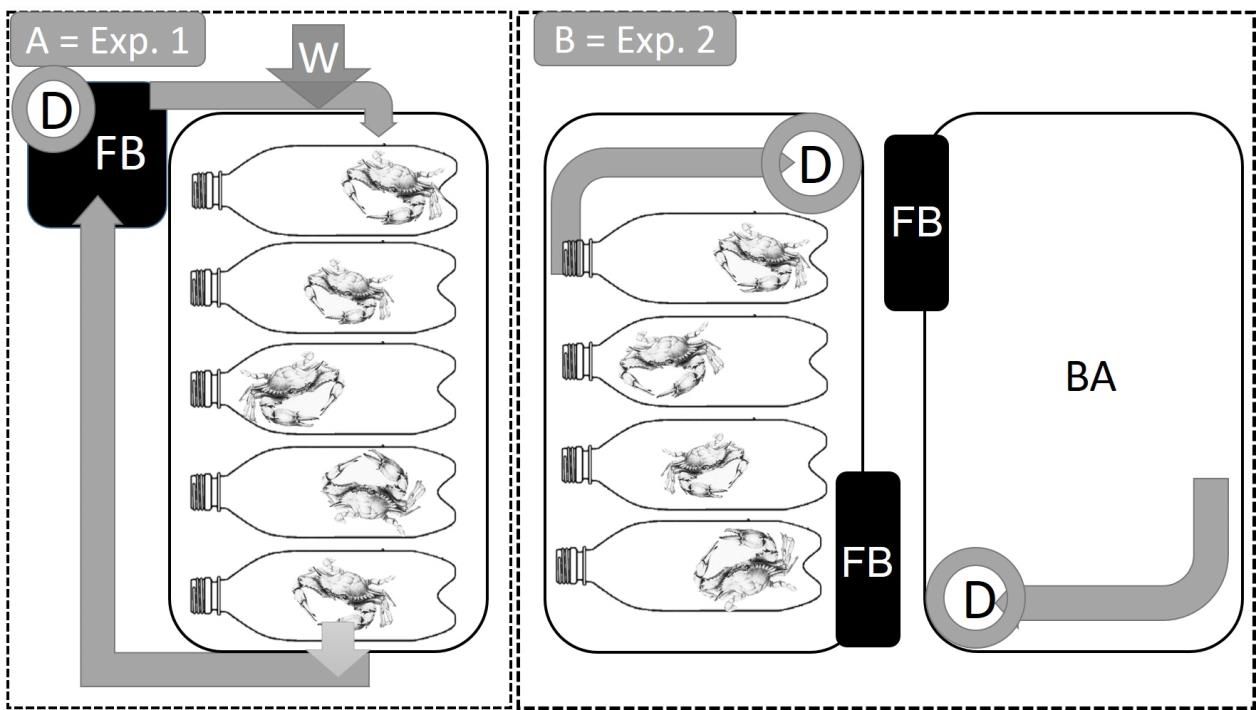
Schematic representations of the experimental systems. A: Experiment 1 (Exp. 1): 66 crabs (46 in pre-ecdysis stage and 20 in inter-ecdysis stage) maintained in a collective system with filtration and partial daily water renewal. B: Experiment 2 (Exp. 2): 175 crabs (123 in pre-ecdysis stage and 52 in inter-ecdysis stage) maintained in a collective system with filtration but without water renewal. A = external water supply; D = protein skimmer; FB = mechanical / biological filter; BA = Water control.

### Experiment 1: **Crab maintenance in a collective system with filtration and partial daily water renewal**

Sixty-six *C. ornatus* crabs were individually placed in perforated pet bottles (600 mL) and distributed in a system consisting of 20 polyethylene tanks (71.0 x 35.5 x 35.0 cm, containing 25 L of seawater each). The tanks were interconnected via a skimmer and a mechanical/biological filter system and were under constant aeration, continuous water recirculation and controlled photoperiod (14L:10D).

The animals were separated into two groups: A1, pre-ecdysis animals (n = 46), and AC, control animals (at the inter-ecdysis stage) (n = 20). Each day throughout the experimental period (12 days), 1/3 of the total water volume of the system (333 L) was added, promoting mixing with the water already present, and an approximately equivalent amount was removed, keeping the total water volume in the system constant.

### Experiment 2: **Crab maintenance in a collective system with filtration and without water renewal**

One hundred and seventy-six *C. ornatus* crabs were individually placed in perforated (600 mL) pet bottles and distributed among 12 polyethylene tanks (71.0 x 35.5 x 35.0 cm, containing approximately 30 L of water each). Each tank contained a protein skimmer and a mechanical/biological filter and was subjected to constant aeration, continuous water recirculation and controlled photoperiod (14L:10D).

The animals were subdivided into 3 groups: B1, pre-ecdysis animals (n = 83); B2, pre-ecdysis animals (n = 40); and BC, control inter-ecdysis animals (n = 52). In addition, three tanks containing water only were maintained throughout the experimental period (12 days) for comparison of physical and chemical water variables between these tanks and the 3 treatment groups. There was no water renewal during the experiment.

The animals in the B2 group were housed in the same tanks used to house group B1 and maintained in the same water used for group B1.

Table 1 provides summary information on the subject animals and design of the two experiments.

**Table 1.** Summary of the general conditions of the experiments performed to evaluate the effects of water quality on the hardening time of the exoskeleton in *Callinectes ornatus* and width and weight data of the animals (mean ± SD). NA: not applicable.

| data of the animals (mean ± SD). N/A: not applicable. | | | | | | | | |
| --- | --- | --- | --- | --- | --- | --- | --- | --- |
| Experiment | Group | Experimental Unit | Stage | N | Width (mm) (±sd) | Weight (g) (±sd) | Water Volume (L) | Water Sampling Frequency |
| 1 | A1 | Tank | Pre-ecdysis | 46 | 45.4 (±6.16) | 15.04 (±6.11) | 0.6 | 24 h |
| | AC | Tank | Inter-ecdysis | 20 | 61.55 (±10.53) | 35.57 (±16.9) | 0.6 | 24 h |
| 2 | B1 | Tank (Replicate 1) | Pre-ecdysis | 27 | 50.7 (±7.78) | 16.64 (±7.03) | 32.5 | 12 h |
| | | Tank (Replicate 2) | | 27 | 3.10 (±7.36) | 14.36 (±7.47) | 32.5 | 12 h |
| | | Tank (Replicate 3) | | 29 | 47.5 (±5.12) | 15.11 (±4.92) | 32.5 | 12 h |
| | B2 | Tank (Replicate 1) | Pre-ecdysis | 14 | 48.1 (±8.25) | 16.46 (±8.08) | 30 | 12 h |
| | | Tank (Replicate 2) | | 13 | 46.4 (±7.02) | 15.40 (±6.7) | 30 | 12 h |
| | | Tank (Replicate 3) | | 13 | 47.0 (±6.64) | 14.75 (±6.3) | 30 | 12 h |
| | BC | Tank (Replicate 1) | Inter-ecdysis | 17 | 56.2 (±2.9) | 26.51 (±5.07) | 32.5 | 12 h |
|  |  | Tank (Replicate 2) |  | 17 | 59.7 | 28.31 | 32.5 | 12 h |

| Experiment | Group | Experimental Unit | Stage | N | Width (mm) (±sd) | Weight (g) (±sd) | Water Volume (L) | Water Sampling Frequency |
| --- | --- | --- | --- | --- | --- | --- | --- | --- |
| | | 2) | | | (±3.8) | (±6.1) | | |
| | | Tank (Replicate 3) | | 18 | 56.65 (±4.93) | 26.54 (±7.95) | 32.5 | 12 h |
|  | BA | Tank (3 Replicates) | NA | NA | NA | NA | 32.5 | 12 h |

### Experimental Procedures

During the experiments, the crabs were monitored every three hours on the first four days, every six hours on the following five days, and every 12 hours on the last three days of experimentation, preferably between 18:00 and 06:00 h. These times were selected based on the results of the pilot experiments.

Monitoring consisted of identifying animals undergoing the moulting process, removing any moulted exoskeletons (to prevent the animals from obtaining calcium by feeding on them), evaluating the consistency of the carapace of those animals that had moulted, and removing any dead animals. Evaluating the consistency of the exoskeleton was performed by pressing the carapace with an index finger. Sufficient pressure was applied to deform the carapace but not injure the animal or break the carapace when rigid. Based on the resistance to pressure and texture of the exoskeleton, its consistency (Co) was classified by the evaluator as follows: hard - before ecdysis (1), soft (2), leather (3), soft paper (4), hard (5) or hard paper - after ecdysis (6). To reduce and standardise the error, a single evaluator performed the consistency assessments in both experiments.

### Water analysis

In both experiments, salinity (refractometer; Instrutemp, Brazil), temperature (digital thermometer), pH (AZ pH/mV/TDS /Temperature Meter 86505, Taiwan), and dissolved oxygen concentration (Oximeter YSI 550A, USA) were monitored daily in all experimental units. Water samples were collected from the units, labelled and immediately frozen (-20 ° C) for later evaluation of the physical and chemical variables. For group A1, water collection was performed every 24 hours before the new water was added to the system. For groups B1, B2, BC and BA, 50 mL of water was collected every 12 h.

At the end of the experiments, the frozen water samples were analysed with respect to the following parameters: Na^+^, K^+^, Ca^2+^ and NO3 (electrodes of the LAQUAtwin series, Horiba Scientific®, Japan) and total ammonia (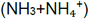) and 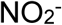 (SpectraMax® m2 spectrophotometer, USA). Measurements were performed following APHA (2005) and Büldt and Karst (1999). The determinations of 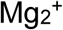and Cl^-^were performed using colourimetry (Labtest®, Brazil) at a wavelength of 540 nm and 470 nm, respectively (SpectraMax® m2, USA), according to the method described by Clarke (1950).

### Statistical analyses

The survival of the animals during the experiments was analysed through Kaplan-Meier curves. The data were grouped by treatment (groups), and the normality of the distribution of each variable was tested by using Shapiro-Wilk test. Where the normality hypothesis was rejected, non-parametric Mann-Whitney or Kruskal-Wallis tests were used.

Multiple linear regression analysis was performed to model the influences of the physical and chemical variables that determine water quality on exoskeleton hardening time. The assumption of the independence of the physical and chemical variables was upheld, the hypothesis of autocorrelation and collinearity (using Durbin-Watson and the serial error correlation tests) was rejected, and the normality of the error was confirmed.

To limit the number of variables and thereby minimise the complexity of the models without a significant loss of the information offered by the total set of original variables, we select only those variables that: 1) were statistically significant (p < 0,05) and; 2) contributed more than 5% to the coefficient of determination (R^2^) of the model or that caused the R^2^ value to move into a higher category when it was included in the model, following the classification proposed by Mukaka (2012): very weak:: R^2^ < 0,19; weak: 0,20 > R^2^ < 0,39; moderate: 0,40 > R^2^ < 0,69; strong: 0,70 > R^2^ < 0,89; very strong: R^2^ > 0,90.

## Results

### Ecdysis

Significant effects of sex on survival rate, ecdysis, or exoskeleton post-ecdysis hardening time were not observed. Therefore, the data from males and females were pooled. In addition, water temperature (27.0 ± 1.1 ° C), salinity (31.0 ± 2.1 ups) and dissolved oxygen concentration (5.0 ± 0.52 mg/L) remained largely stable and did not significantly influence any of the dependent variables.

The moulting rate of the animals in pre-ecdysis at the beginning of the experiments ranged from 40 to 95%. Most moulting events occurred during the night, and 50% of the animals moulted between 52 and 80 hours after the beginning of the experiments. There was a significant effect of moulting on final mortality rate and on survival time after ecdysis (Table 2).

**Table 2.**
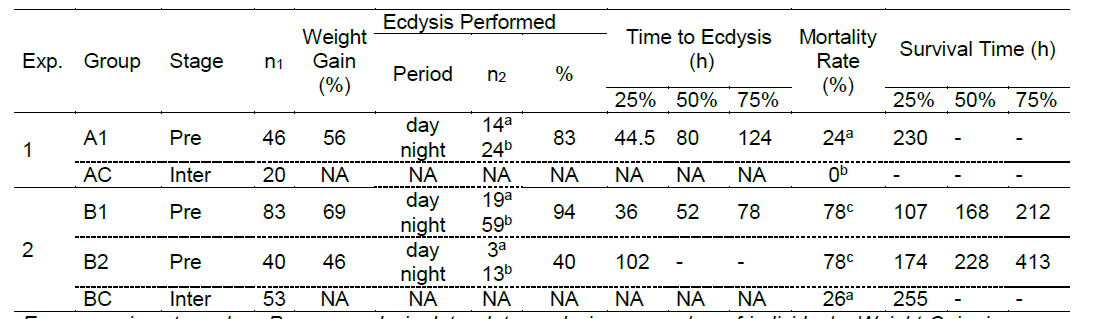
General results of laboratory experiments to evaluate ecdysis in *Callinectes ornatus.*

*Exp.: experiment number; Pre: pre-ecdysis; Inter: Inter-ecdysis; n1: number of individuals; Weight Gain: increase in post-ecdysis weight (%); Period: the period in which moult occurred; n2: number and percentage of crabs that performed ecdysis; Time to Ecdysis: time (h) at which 25, 50 and 75% of the animals had moulted; Mortality rate (%);Survival Time: time (h) at which 25, 50 and 75% of the animals survived after ecdysis; NA: Not Applicable. Different letters indicate significant differences (p < 0.05) between the groups according to the Kruskal-Wallis test. Experiment 1 (collective treatment with filtration and partial daily water renovation). A1: pre-ecdysis organisms; AC (Control): organisms in inter-ecdysis. Experiment 2 (collective treatment with filtration but no water renewal). B1: pre-ecdysis organisms; B2: tanks containing water previously used for group B1, with organisms in pre-ecdysis; BC (Control): tanks with organisms in inter-ecdysis.*

### Physical and chemical water parameters

In Experiment 1, the total ammonia (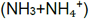) concentrations remained below the limit of analytical detection, and the median pH was 8.5, with variation between 8.1 and 8.5. The remaining physical and chemical parameters were relatively stable throughout the experimental period (Table 3).

**Table 3.** Median and 1st and 3rd quartiles of the water quality parameters in Experiment 1 (collective treatment with filtration and partial daily water renewal).

| Parameter | Median | 25-75% |
| --- | --- | --- |
| pH | 8.50 | 8.4-8.5 |
| K<sup>+</sup> (mg/L) | 380 | 370-390 |
| Ca<sup>2+</sup> (mg/L) | 350 | 330-430 |
| Mg<sup>2+</sup> (mg/L) | 589.5 | 573.3-602.6 |
| Na<sup>+</sup> (mg/L) | 11,000 | 9,900-12,000 |
| Cl<sup>-</sup> (mg/L) | 16,830 | 16,059-17,668 |
| TA (mg/L) | 0.0 | 0.0 |
| NH<sub>3</sub> (mg/L) | 0.0 | 0.0 |
| NO<sub>2</sub><sup>-</sup> (mg/L) | 1.30 | 1.29-1.30 |
| NO<sub>3</sub> (mg/L) | 180 | 170-200 |

TA: Total ammonia (NH<sub>3</sub>+NH<sub>4</sub><sup>+</sup>)

In Experiment 2, only potassium and sodium concentrations presented differences between the groups B1 and B2. There was a reduction in pH and increases in total ammonia and nitrite concentrations in the experimental treatments (pre-ecdysis organisms, B1 and B2) in relation to the control (BA, tanks containing water only). The variables monitored in the BC tanks (inter-ecdysis organisms) presented intermediate values relative to the other groups (Table 4).

**Table 4.** Median and 1st and 3rd quartiles of the water parameters in Experiment 2 (collective treatment with filtration but without water renewal). *B1: organisms in pre-ecdysis. B2: tanks containing water previously used for group B1 and pre-ecdysis crabs. BA (Control): tanks containing water only. BC (Control): tanks with crabs in inter-ecdysis. Different letters indicate significant differences (p < 0.05) between groups according to the Kruskal-Wallis test. TA: Total ammonia (NH3+NH4+).*

| Parameter | Groups |  |  |  |
| --- | --- | --- | --- | --- |
|  | B1 | B2 | BA | BC |
|  | Median<br>25-75% | Median<br>25-75% | Median<br>25-75% | Median<br>25-75% |
| pH | 6.7<sup>b</sup><br>(6.3-7) | 6.5<sup>b</sup><br>(6.2-7) | 8.4<sup>a</sup><br>(8.3-8.4) | 7.8<sup>ab</sup><br>(6.8-8.1) |
| K<sup>-</sup> (mg/L) | 420.00<sup>a</sup><br>(370-540) | 280.00<sup>b</sup><br>(230-330) | 330.00<sup>ab</sup><br>(260-370) | 390.00<sup>ab</sup><br>(290-420) |
| Ca<sup>2+</sup> (mg/L) | 430.00<sup>a</sup><br>(400-480) | 350.00<sup>a</sup><br>(320-420) | 430.00<sup>a</sup><br>(360-490) | 450.00<sup>a</sup><br>(390-480) |
| Mg<sup>2+</sup> (mg/L) | 545.6<sup>b</sup><br>(472.3-585.8) | 551.2<sup>b</sup><br>(531.8-622.5) | 586.3<sup>ab</sup><br>(573.3-597.4) | 592.0<sup>a</sup><br>(555.2-607.9) |
| Na<sup>+</sup> (mg/L) | 12,000<sup>a</sup><br>(10,000-14,000) | 7,500<sup>b</sup><br>(6,500-9,000) | 11,500<sup>ab</sup><br>(10,000-13,000) | 12,000<sup>ab</sup><br>(10,000-14,000) |
| Cl<sup>-</sup> (mg/L) | 18279<sup>a</sup><br>(16,507-20,245) | 11995<sup>a</sup><br>(9,158-15,782) | 15830<sup>a</sup><br>(14,705-17,473) | 17602<sup>a</sup><br>(16,122-19,278) |
| TA (mg/L) | 6.8<sup>b</sup><br>(5.7-10.5) | 9.5<sup>b</sup><br>(7.4-11.91) | 0.0<sup>a</sup><br>0.0 | 1.1<sup>ab</sup><br>(0.0-3.1) |
| NH<sub>3</sub> (mg/L) | 0.02<sup>b</sup><br>(0.01-0.05) | 0.02<sup>b</sup><br>(0.01-0.05) | 0.00<sup>a</sup><br>0 | 0.01<sup>ab</sup><br>(0-0.08) |
| NO<sub>2</sub><sup>-</sup> (mg/L) | 4.7<sup>b</sup><br>(2.8-6.5) | 5.4<sup>b</sup><br>(2.15-7.8) | 1.3<sup>a</sup><br>(1.3-1.4) | 5.2<sup>b</sup><br>(4.0-6.1) |
| NO<sub>3</sub> (mg/L) | 230<sup>a</sup><br>(190-310) | 260<sup>a</sup><br>(200-340) | 210<sup>a</sup><br>(120-260) | 230<sup>a</sup><br>(150-280) |

### Influence of the physical and chemical water parameters on the survival and moulting of *C. ornatus*

As expected, crab survival time was influenced by moulting regardless of the experiment. The organisms of group B1 that underwent moult in the first 36 h showed rapid exoskeleton hardening and low mortality rates. Therefore, the B1 data were divided into two categories: M1, animals that moulted within the first 36 h, and M2, those that moulted after 36 h. The survival of M1 animals was strongly influenced by pH and total ammonia and nitrite concentrations, whereas the survival of the M2 animals was moderately influenced by the same variables.

Table 5 shows the multiple linear regression results. Crab survival rate was significantly influenced by pH, nitrite and total ammonia in all of the experiments. The remaining parameters had no significant influence (p < 0.05) on the crab survival time.

**Table 5.**
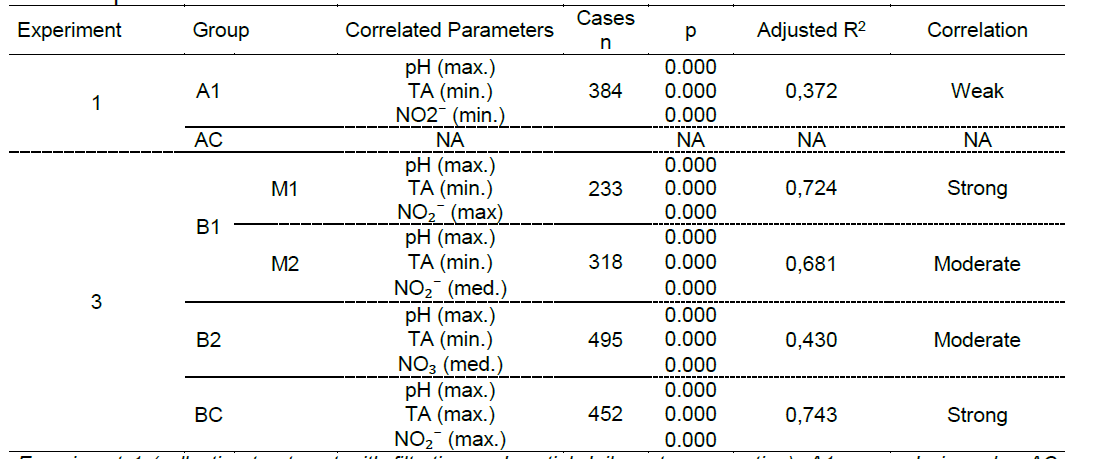
General results of multiple linear regression analysis of the influences of physical and chemical parameters on crab survival time. *Experiment 1 (collective treatment with filtration and partial daily water renovation). A1: pre-ecdysis crabs; AC (Control): inter-ecdysis crabs. Experiment 2: Collective treatment with filtration but no water renewal. B1: pre-ecdysis crabs; B2: tanks containing water previously used for the B1 group, with pre-ecdysis crabs; BC (Control): tanks with crabs in inter-ecdysis; M1: crabs of B1 Group that performed ecdysis within the first 36 hours; M2: crabs that moulted after 36 hours. NA: Not applicable. TA: Total ammonia (NH3+NH4+).*

As expected, crab survival time was influenced by moulting regardless of the experiment. The organisms of group B1 that underwent moult in the first 36 h showed rapid exoskeleton hardening and low mortality rates. Therefore, the B1 data were divided into two categories: M1, animals that moulted within the first 36 h, and M2, those that moulted after 36 h. The survival of M1 animals was strongly influenced by pH and total ammonia and nitrite concentrations, whereas the survival of the M2 animals was moderately influenced by the same variables.

The results of the multiple linear regression analysis of the effects of the physical and chemical parameters on the time until either the shells fully hardened (reached Co 6) or death are presented in Table 6. In Experiment 1, only pH had an influence (weak) on the results. In experiment 2, pH, ammonia and nitrite had moderate influences on the results.

**Table 6.**
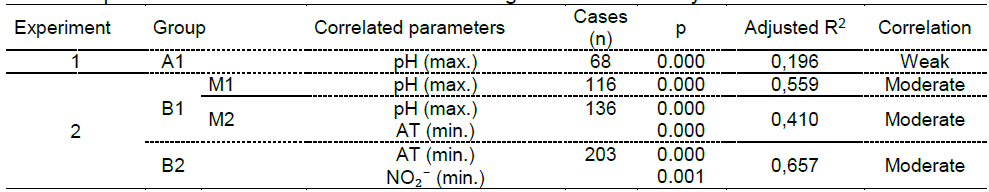
General results of multiple linear regression analysis of the influences of physical and chemical parameters on the time until shell hardening or death after ecdysis. *Experiment 1 (collective treatment with filtration and partial daily water renovation). A1: crabs in pre-ecdysis. Experiment 2 (collective treatment with filtration but no water renewal). B1: crabs in pre-ecdysis; B2: tanks containing water previously used for the B1 group, with crabs also in pre-ecdysis; M1: crabs that moulted within 36 hours; M2: crabs that moulted after 36 hours. TA: Total ammonia (NH3+NH4+).*

The duration at which the shell was at consistency 2 (i.e., the consistency with the highest market value) was significantly higher in the M2 animals than in the M1 animals. Furthermore, none of the M2 individuals that moulted after 36 h achieved Co 6 (hard) shells, whereas in the M1 group, more than half of the individuals had shells that reached this consistency. In addition, 68% of individuals with shells that hardened remain alive. Among those that did not achieve shell hardening, the survival rate was only 15% (Table 7).

**Table 7.**
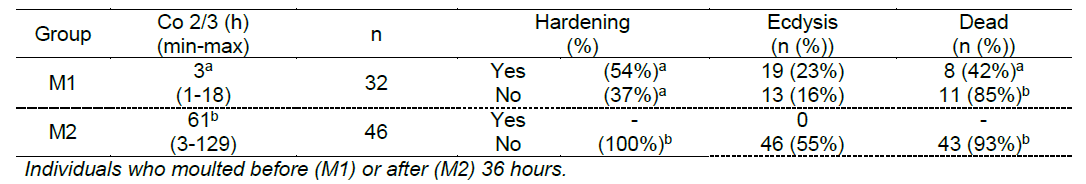
Duration of *Callinectes ornatus* at shell consistencies 2 and 3 (Co 2 and Co 3) and associated hardening, ecdysis and mortality data in group B1 (organisms initially in pre-ecdysis) of Experiment 2 (collective treatment with filtration but no water renewal). Different letters indicate a significant difference (p < 0.05) between the groups (within a column) according to the Kruskal-Wallis or Mann-Whitney test.

Table 8 shows the median, minimum and maximum values of the physical and chemical parameters of water quality that most influenced crab survival time and exoskeleton consistency: pH, ammonia and nitrite. The duration at each level of consistency was highly related to the pH and concentration of total ammonia at the time of moulting. When the levels of the physical and chemical parameters favoured hardening, the time until the organisms reached Co 4 (soft paper) was short (between two and four hours). In contrast, when the pH was below 7.3 and the total ammonia concentration remained above 6 mg/L, the median time to Co 4 was 60 h.

**Table 8.**
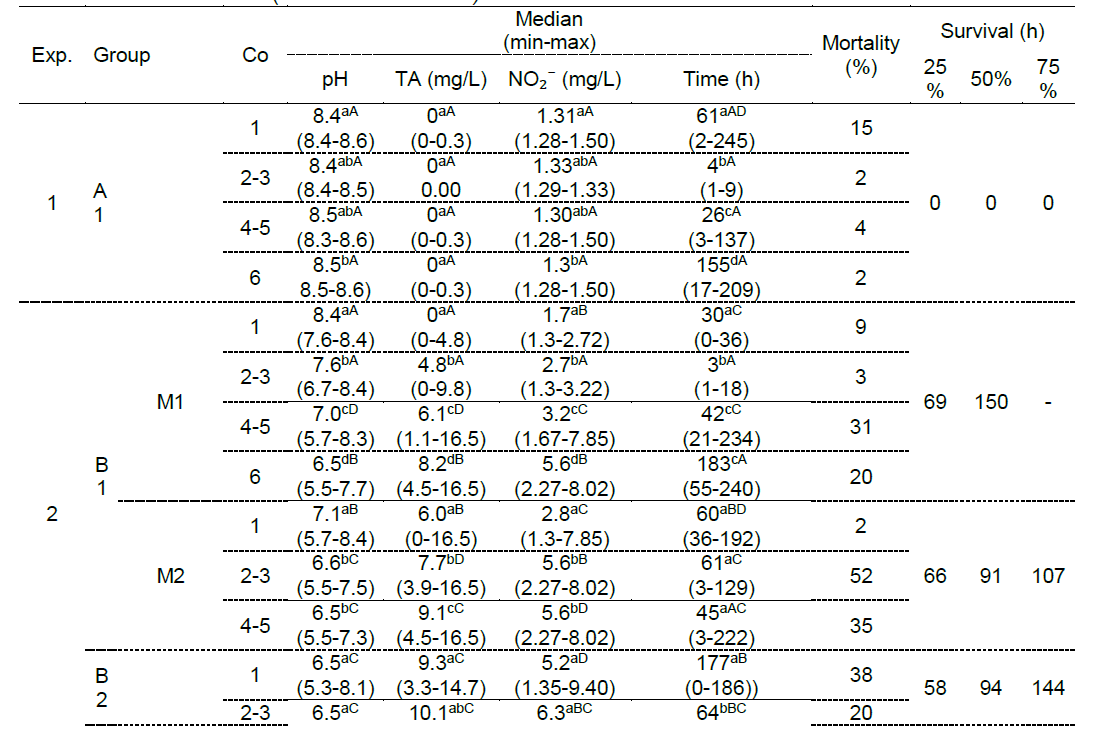
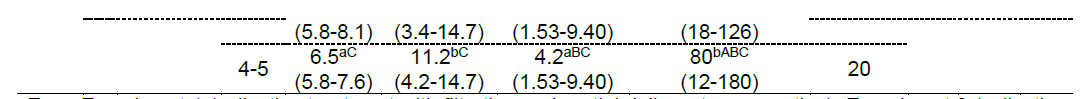
Water quality parameter measurements and crab survival data registered in the experimental units in which the crabs (*Callinectes ornatus)* moulted. *Exp.: Experiment 1 (collective treatment with filtration and partial daily water renovation), Experiment 2 (collective treatment with filtration but no water renewal). A1 and B1: water not used previously; B2: tanks containing water used previously for the group B1; M1: crabs that moulted within the first 36 hours; M2: crabs that moulted after 36 hours. Co: consistency. TA: total ammonia. Time: Length of stay at a given consistency. Different letters indicate significant differences (p < 0.05) according to the Kruskal-Wallis test. Lowercase letters indicate differences in carapace consistency within the same group. Uppercase letters indicate differences in carapace consistency among groups.*

When water renewal was not performed (Experiment 2), the pH and total ammonia and nitrite concentrations varied significantly (Figure 2). As a result, there was an increase in the carapace hardening time and a decrease in the number of individuals reaching Co 6. The crabs of Experiment 1 (A1) and the animals that moulted within the first 36 hours of Experiment 2 (M1) spent significantly less time at Co 2 and Co 3 than did those that moulted after the first 36 hours (M2) in group B1 and those in group B2.

**Figure 2.**
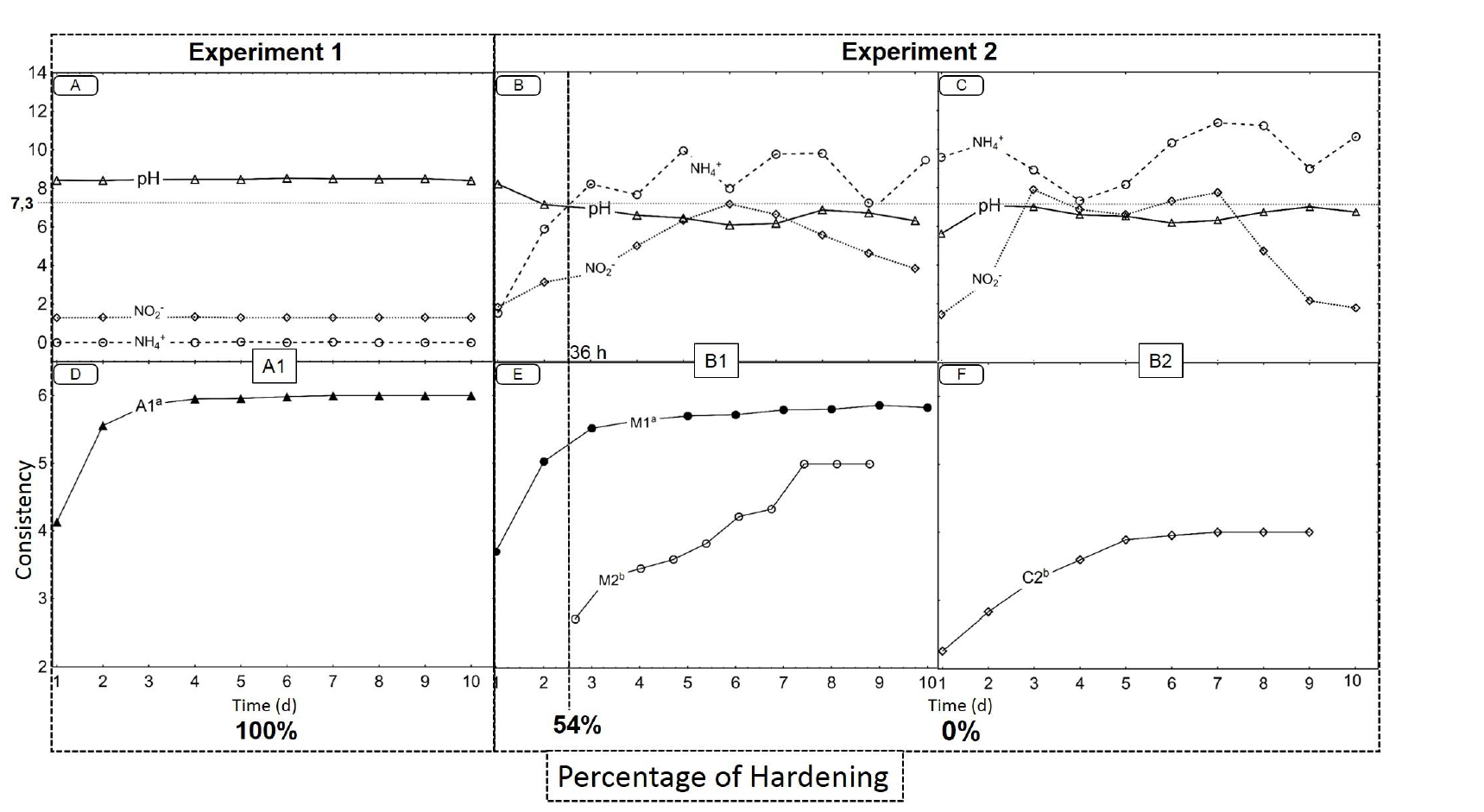
Median pH, total ammonia (mg/L), nitrite (NO2^-^) (mg/L) (A, B and C) and carapace consistency (2 - soft, 3-leather, 4 - soft paper, 5 - hard, paper 6 - hard) over time (in days) (D, E and F). Experiment 1: collective treatment with filtration and partial daily water renewal (66 crabs tested, 46 in pre-ecdysis and 20 in inter-ecdysis stage). Experiment 2: collective treatment with filtration but without water renewal (175 crabs tested, 123 in pre-ecdysis and 52 in inter-ecdysis stage). A1 (46 crabs in pre-ecdysis stage) and B1 (86 crabs in pre-ecdysis stage divided in three replicas): previously unused water and organisms in pre-ecdysis. B2 (40 crabs in pre-ecdysis stage): organisms maintained in the reused water of group B1. M1: 32 crabs that moulted within the first 36 hours. M2: 46 crabs that moulted after 36 hours. Different letters indicate significant differences (p < 0.05) among groups according to the Kruskal-Wallis test.

## Discussion

An issue repeatedly debated among those who investigate the shedding and hardening process in crustaceans is the importance of calcium, the main constituent element of the exoskeleton (Greenaway, 1985), in this process (Cameron, 1985; Cameron and Wood, 1985; Clarke and Wheeler, 1922; Freeman and Perry, 1985; Granado e Sá et al., 2010; Greenaway, 1983; Mangum et al., 1985; Middlemiss et al., 2016; Neufeld and Cameron, 1992; Pan et al., 2006; Perry et al., 2001; Robertson, 1960; Welinder, 1974; Wheatly et al., 2001; Wheatly, 1997; Wheatly, 1999; Wheatly et al., 2002; Zanotto and Wheatly, 2002). The lack of significant correlations between the concentrations of Ca^2+^ and Mg^2+^ in water and either carapace hardening or *C. ornatus* survival does not indicate that calcium is not important in this process. On the contrary, it indicates that certain processes can directly interfere with the physiology of the absorption and immobilisation of Ca2^+^ in the exoskeleton and thereby significantly increase the time that these animals remain soft after moult.

The organisms of Experiment 1 (subjected to daily water renewal) that underwent ecdysis hardened rapidly, achieving paper consistency (Co 4) a median of 4 h after moult. This finding is consistent with studies conducted with *C. sapidus* (Cameron and Wood, 1985; Freeman et al., 1987). Similar results were observed among the crabs in Experiment 2 that moulted in water with a pH above 7.6 and a total ammonia concentration below 4.8 mg/L, with Co 4 achieved after a median time of 2 to 3 h. However, among the animals that began moulting in water with a pH below 7.3 and a total ammonia concentration above 6 mg/L, up to 129 h (median of more than 60 h) elapsed before either reaching Co 4 or death.

To understand this result, it is necessary to understand the chemical processes involved in the calcification of the crab exoskeleton. In a closed system with water recirculation, it is expected that over time there will be a reduction in the concentration of free Ca2^+^, due mainly to the immobilisation of Ca2^+^ in the form of CaCO3 during exoskeleton hardening (Perry et al., 2001). This immobilisation can be represented by the following equation:

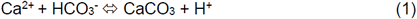

With the increased demands for Ca2^+^ and HCO3^-^, crabs begin to consume both metabolic and external CO2. CO2 reaches its highest internal concentrations at moulting time (Mangum et al., 1985), increasing the availability of internal HCO3^-^ (Cameron and Wood, 1985). As soon as moulting occurs, the enzyme carbonic anhydrase (CA), present mainly in the epithelium and the gills, is activated (Mangum et al., 1985), accelerating the reaction:

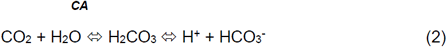

As explained by Detours et al. (1968) and Zeebe and Wolf-Gladrow (2001), the formed carbonic acid tends to be buffered by the carbonate-bicarbonate system. This process results in an increase in the fraction of CO3^-^ and acidification of the medium (Greenaway, 1974; Mangum et al., 1985; Wheatly, 1997). However, over time, the natural acid neutralisation capacity of the system becomes compromised, and the medium tends to acidify as a result, increasingly compromising the crab's capacity to deposit CaCO3 in its exoskeleton. According to Cameron and Wood (1985), the calcification process can be compromised if the pH outside the body is less than 0.3 to 0.5 above the internal pH.

However, in addition to consuming HCO3^-^ post-ecdysis, the organism excretes H^+^ or an equivalent ion such as NH4^+^ (Cameron, 1985; Middlemiss et al., 2016), which is dissociated into NH3 and H^+^. The rate of H^+^/NH4^+^ excretion increases after moulting (Cameron and Wood, 1985) and may increase further during bacterial denitrification (Rijn et al., 2005). Under these conditions, the metabolism of excretion also contributes to the acidification of the medium, further reducing the capacity for calcium mobilisation by the crab, as observed in experiment 2. There is evidence that water acidification is more critical for the hardening process of marine crustaceans than for that of freshwater crustaceans. Unlike freshwater crustaceans, marine crustaceans have almost no internal reserves of calcium (gastroliths) and depend exclusively on the environment to supply the demand for Ca^2+^ (Greenaway, 1985; Passano, 1960; Wheatly, 1997).

In a similar manner, acidification might affect the deposition of magnesium in the crustacean exoskeleton. Although magnesium concentrations in water are relatively lower than those of calcium, magnesium also plays an important role in the hardening of the exoskeleton, and it is also obtained through water (Cameron and Wood, 1985; Clarke and Wheeler, 1922; Welinder, 1974) in a process that might be affected by pH (Tao et al., 2009).

In addition to Ca^2+^ and Mg^2+^ concentrations, the concentrations of Na^+^ and K^+^ were monitored in this study. These two ions directly participate in important enzymatic activities that occur post-ecdysis (Towle and Mangum, 1985). Studies have shown that if the relative proportions of these two ions are altered, ammonia toxicity can occur due to the retention of ammonia by the organism and potentially compromise the anima's survival (Pan et al., 2006; Romano and Zeng, 2011; Zanotto and Wheatly, 1993). However, were observed no significant effects of these ions in our experiments. It is possible that the factors described above were much more important in influencing exoskeleton hardening and the probability of survival in *C. ornatus*.

There was also a direct relationship between the time to exoskeleton hardening and the mortality rate. However, the mortality rate was only 25% among those crabs that moulted after approximately 60 h. Those that did not moult died or remained alive until the end of the experiment. In addition, in all of the groups except those receiving periodic water renewal, there was an increase in mortality in the post-ecdysis phase. In this case, the analyses again indicated the influences of pH and total ammonia.

It is known that crabs (notably *C. sapidus*, the most studied species of the genus *Callinectes*) can tolerate a pH range of 6.5 to 8.5 (Hochheimer, 1988). Nevertheless, in artificial environments, it is recommended that pH be maintained between 7.0 and 8.0 (Oesterling, 1995). It is also known that there are behavioural and tolerance differences between young and adult animals in relation to pH (Laughlin et al., 1978). In Experiment 2 of the present study, pH values of 5.5 and 5.3 were recorded in groups B1 and B2, respectively. In addition to having a direct effect on the organisms, a reduction of pH causes an increase in the nitrous acid fraction (HNO2) present in water; HNO2 is toxic to aquatic organisms (Ary and Poirrier, 1989; Lin and Chen, 2003; Russo et al., 1981; Seneriches-Abiera et al., 2007).

The toxicity of ammonia, in turn, is directly proportional to pH and NH3 concentrations. Romano and Zeng (2007) estimated an LC50 for juveniles of *Scylla serrata* of 6.81 mg/L NH3-N. Koo et al. (2005) *reported that at least 50% of juveniles of Orithyia sinica* survived for 30 days at approximately 2.33 mg/L NH3-N. Lakshmi (1984) reported a mortality rate of 20% in *C. sapidus* in pre-ecdysis at 1.41 mg/L NH3, which increased to 100% at 2.31 mg/L NH3. In our experiments, a pH reduction was observed over time, which indicated that the NH3 concentrations remained sufficiently low as to rule out any toxic effects of ammonia on *C. ornatus*.

Regarding nitrite, there is no consensus regarding the concentrations at which this compound is toxic to crabs. Lakshmi (1984) and Ary and Poirrier (1989) reported that the survival of *C. sapidus* was only affected at NO2^-^ concentrations above 10 mg/L. According to those authors, crab mortality reached 100% only after 96 h of exposure to concentrations between 50 and 150 mg/L in water with a pH close to 8. In contrast, Manthe et al. (1984) found that the moulting efficiency of *C. sapidus* was affected by nitrite concentrations close to 2 mg/L. In the present study, the nitrite concentration reached 7.6 mg/L. Thus, it is possible that the observed mortality might have been influenced by both pH and nitrite levels during the experiments and that they had a cumulative effect. Moreover, a long hardening time, which exposed the animals to unfavourable physiological conditions, appears to have significantly increased the risk of death.

## Conclusion

Moulting in *C. ornatus* exhibited strong relationships with the characteristics of the crab's aquatic medium. The crabs drastically altered the physical and chemical characteristics of the water, mainly through processes related to acidification and ammonification. These alterations, in turn, directly interfered with exoskeleton hardening, causing the exoskeletons of the animals to remain at soft or paper consistency for periods of up to 5 days. Commercially, the establishment of such periods would allow crabs to be marketed as soft-shell crabs within a time window more than 20 times longer than that typically observed. If the results observed here can be replicated at the commercial scale, large reductions in workload and operational costs could be obtained, increasing the efficiency and viability of large-scale crab production.

## Acknowledgements

We thank GIA (Grupo Integrado de Aquicultura e Estudos Ambientais), Universidade Federal do Paraná and Instituto Federal do Paraná for the opportunity of conducting soft-crabs studies. Thanks to Vitor Rossi, Marcelo F. A. Pinto, Nathiele Cozer and Amanda C. Alburquerque for the collaboration during the experiments. Additionally, we thank the fishermen Zeca Tavares, Túlio and Andréa for the crabs experimental capture. This study is a result of the Master degree thesis of DBH under the post-graduate initiative “PPG Zootecnia, Universidade Federal do Paraná - UFPR”.

## Funding

This study was supported by the “Conselho Nacional de Desenvolvimento Científico e Tecnológico – CNPq” (process 302609/2013-0) granted to Dr Antonio Ostrensky, and by CNPq financing of the projects associated with this manuscript (processes 381091/2014-7, 473959-2013, 403705/2013-4 and 468251/2014-6). DBH receives a scholarship from CNPq (DTI, process 381091/2014-7). ACR receives a scholarship from CAPES Pró-Amazônia Program (process 1644571).

